# *In vitro* evaluation of the response of human tendon-derived stromal cells to a novel electrospun suture for tendon repair

**DOI:** 10.1101/2020.11.11.378695

**Authors:** A Nezhentsev, RE Abhari, MJ Baldwin, JY Mimpen, E Augustyniak, M Isaacs, P-A Mouthuy, AJ Carr, SJB Snelling

## Abstract

Recurrent tears after surgical tendon repair remain common. Repair failures can be partly attributed to the use of sutures not designed for the tendon cellular niche nor for the promotion of repair processes. Synthetic electrospun materials can mechanically support the tendon whilst providing topographical cues that regulate cell behaviour. Here, a novel electrospun suture made from twisted polydioxanone (PDO) polymer filaments is compared to PDS II, a clinically-used PDO suture currently utilised in tendon repair. We evaluated the ability of these sutures to support the attachment and proliferation of human tendon-derived stromal cells using PrestoBlue and Scanning Electron Microscopy. Suture surface chemistry was analysed using X-ray Photoelectron Spectroscopy. Bulk RNA-Seq interrogated the transcriptional response of primary tendon-derived stromal cells to sutures after 14 days. Electrospun suture showed increased initial cell attachment and a stronger transcriptional response compared to PDS II, with relative enrichment of pathways including mTorc1 signalling and depletion of epithelial mesenchymal transition. Neither suture induced transcriptional upregulation of inflammatory pathways compared to baseline. Twisted electrospun sutures therefore show promise in improving outcomes in surgical tendon repair by allowing increased cell attachment whilst maintaining an appropriate tissue response.

## Introduction

Tendon injuries are common and cause pain and reduced quality of life for patients. Rotator cuff tendon tears alone affect around 50% of those over 66 years of age^1^ and many of these patients require surgical repair^2^. However, despite exploration of different suture methods, surgical tendon repairs have poor outcomes due to suture pull-through or tissue re-tears, with 40% of rotator cuff repairs failing within one year^3^. This failure to successfully repair torn tendons causes long-term disability and represents a significant socioeconomic cost^4^.

Suture-based re-joining of tendon ends is the current gold standard surgical treatment for tendon injuries, but currently used sutures have been repurposed from other anatomical sites and have not been designed for the tendon niche. Clinically-used sutures are manufactured from synthetic polymers with high tensile strength to provide mechanical augmentation. However, their topographical characteristics (including fibre diameter) are dissimilar to native tendon, which is composed of multiple aligned small diameter collagen fibres. Furthermore, cells do not integrate with the material^5^, demonstrated by acellular zones forming around sutured sites^6,7^. This lack of infiltration and integration likely contributes to poor tendon healing at the suture-tissue interface. In addition, immune and stromal cells drive a foreign body response to implantable materials including sutures. This can result in chronic inflammation, which can hamper endogenous repair and lead to tissue failure^8^.

The use of sutures that enable cellular attachment and proliferation, and that do not raise a chronic inflammatory response may improve the efficacy of tendon repair^5^. Electrospun materials show particular promise and can be produced from synthetic and clinically-approved materials, including polydioxanone (PDO), which is an absorbable polymer used to produce PDS II sutures currently deployed in tendon repair. The high surface area and porosity of electrospun materials together with the ability to manufacture them with fibre diameters similar to the collagen fibres in native tendons provides biomimicry not afforded by conventional sutures^9^. Electrospun materials also promote infiltration and proliferation of stromal cells, including those derived from tendon, and induce expression of tenogenic markers^10–12^. Surface chemistry, fibre diameter, and alignment of materials can tune cell behaviour and this can be exploited to drive repair. However, when manipulating the mechanical and topographical properties of a clinically-approved material it is important to define the cellular response to its specific material properties.

Twisting and braiding electrospun strands into bundles can direct cells to attach and grow in a parallel network, similar to the macroscopic architecture of tendon^14,15^. The morphology and proliferation of tendon-derived stromal cells cultured on electrospun materials have been previously described, but only a small number of selected genes was assessed in tendon-derived stromal cell gene expression analyses^14,16–18^ and response^7,13^. Previous work has also described the twisting of electrospun PDO fibres into prototype multifilament yarns that have similar a tensile strength to currently used sutures and that resemble the hierarchical structure of tendons^13^. Following *in vivo* tendon injury in a rat and sheep model, prototype electrospun sutures supported cellular infiltration with a minimal inflammatory response. Understanding how tendon cells interact with the material *in vitro* can therefore help predict how the electrospun suture will interact and integrate with tendon *in vivo*. The effect of twisted electrospun sutures on the global transcriptional profile of tendon-derived stromal cells *in vitro* or *in vivo* is not well explored. It therefore remains necessary to understand the response of tendon cells to sutures developed for their surgical repair, compared to sutures currently used in surgery.

The overarching aim of this *in vitro* study is to assess the potential of twisted electrospun PDO sutures in tendon repair. This builds on our previously published work^13,19^ and uses a modified electrospun suture that ensures uniformity of fibre diameter, and a high tensile strength and hierarchical structure that has potential to both mechanically and biologically support repair. We hypothesized that a twisted electrospun suture would promote tendon-derived stromal cell attachment and proliferation and induce a pro-reparative gene expression profile.

## Materials and Methods

### Suture preparation

Electrospun sutures were fabricated according to the protocol described in supplementary material 1. Four 2 cm pieces of electrospun suture and PDS II (Ethicon Inc.) were then melted together at both ends by holding near a 200°C hot wire (Proxxon, Axminster, UK). This formed mats that could be transferred between tissue culture wells without disrupting cells. The mats were sterilised by submerging in 70% ethanol for 2 hours, and dried overnight. The sutures were washed twice in PBS and soaked for 2 hours in D10 medium (DMEM-F12 (Thermo Fisher Scientific) supplemented with 10% Foetal Calf Serum (Labtech, Melbourn, UK) and 1% Penicillin-Streptomycin (Thermo Fisher Scientific)).

### Tendon-derived stromal cell seeding

Waste healthy mid-body hamstring tendon tissue was collected from four male patients aged 20-43 (SD +/− 9.4 yr), BMI 21.46-27.76 (SD +/− 2.77 kg/m2), during Anterior Cruciate Ligament reconstruction. Tendon-derived stromal cells were extracted and expanded from tendon explants as previously described^18^ (S2). 50,000 tendon-derived stromal cells (passage 3) were seeded dropwise onto the suture mats, or directly into empty wells for Tissue Culture Plastic (TCP) controls (3 technical repeats for each patient). TCP controls were used later to determine whether changes in gene expression were caused by cell-instructive cues provided by the suture materials. After 4 hours, mats were transferred into a new 12-well plate.

### Scanning Electron Microscopy

Suture pieces were prepared for imaging (S3). The Evo LS15 Variable Pressure Scanning Electron Microscope (Carl Zeiss AG, Oberkochen, Germany) was used to capture images. Images were taken at 200X magnification to visualise suture structure, and at 2000X magnification to visualise attached cells.

### Cell Viability

Measurements of cell viability were performed by incubating cells seeded on TCP and sutures in 10% PrestoBlue for 2 hours on days 1, 4, 7, 11 and 13 after seeding (S4). Fluorescence was measured using a FLUOstar Omega Microplate Reader (BMG Labtech, Aylesbury, UK) at 544 nm excitation and 590 nm emission.

### RNA Sequencing

RNA was extracted (S5) from hamstring tendon-derived stromal cells (n=4 patients) after 14 days culture on sutures and TCP after 14 days culture and at baseline (time of seeding on sutures and TCP).^20^

Bulk RNA-Seq interrogated the tendon-derived stromal cells’ transcriptomic response to the sutures. Libraries were created using a NEBNext Ultra II Directional RNA Library Prep Kit for Illumina (New England Biolabs, Ipswich, MA, USA) and sequencing performed on an Illumina NextSeq 500 using a NextSeq High Output Kit. A quality control (QC) report was collated using the MultiQC tool (v1.7)^21^. Due to low percentage alignment with the reference genome, one PDS II sample did not pass QC and was excluded from analysis. Principal Component Analysis plots extracted the components responsible for most of the dataset variance (S6). The DESeq2 package^22^ was utilised to undertake pairwise comparisons of gene expression at 14 days of culture. Gene-set enrichment analyses was performed using the cluster Profiler package^23^, and data visualised using Enhanced Volcano and ggplot2.

### X-ray Photoelectron Spectroscopy

To assess the surface chemistry of sutures, sutures were flattened into sheets by a hydraulic press (Specac, Orpington, UK) set to 8 tonnes for 30 seconds, and mounted onto a glass slide using double-sided carbon tape. X-ray Photoelectron Spectroscopy (XPS) measurements were made using an AXIS Supra (KRATOS Analytical, Stretford, UK) (S8).

### Data Analysis

GraphPad Prism 8 (GraphPad Software Inc., San Diego, CA, USA) was used for statistical analysis of XPS and cell viability data. The D’Agostino-Pearson test was used to test the normality of the biological replicates. Unpaired t-tests were used to determine whether there was a difference in between the two suture samples. The Holm-Sidak method was used to correct for multiple comparisons. Results were deemed statistically significant when p<0.05, and statistical significance is displayed in shorthand as *p<0.05, **p<0.01, ***p<0.001, and ****p<0.0001. R and R packages (S7) were used to analyse gene expression data.

## Results

### Initial cell attachment was higher on twisted electrospun suture than PDS II

To compare the ability of electrospun and PDS II sutures to support attachment and proliferation of tendon-derived stromal cells, cells were cultured for 13 days on each surface and assessed using SEM imaging and PrestoBlue viability assays. SEM images showed cells attaching to and proliferating on both sutures, with cell coverage increasing throughout the duration of the experiment (Figure 1 A-H). PrestoBlue viability analysis was used to give a surrogate measure of relative cell number over 13 days of tissue culture. The number of viable cells was higher on the electrospun suture compared to PDS II on day 1 (p<0.001) (Figure 1I), indicating higher initial stromal cell attachment. The number of cells present on electrospun sutures remained higher throughout the culture period, with the electrospun suture containing 4 times as many cells than PDS II by day 13 (p<0.0001) (Figure 1I). When cell number was baseline-corrected relative to day 1, there were no differences in proliferation rates on each suture material (Figure 1J).

**Figure 1:**
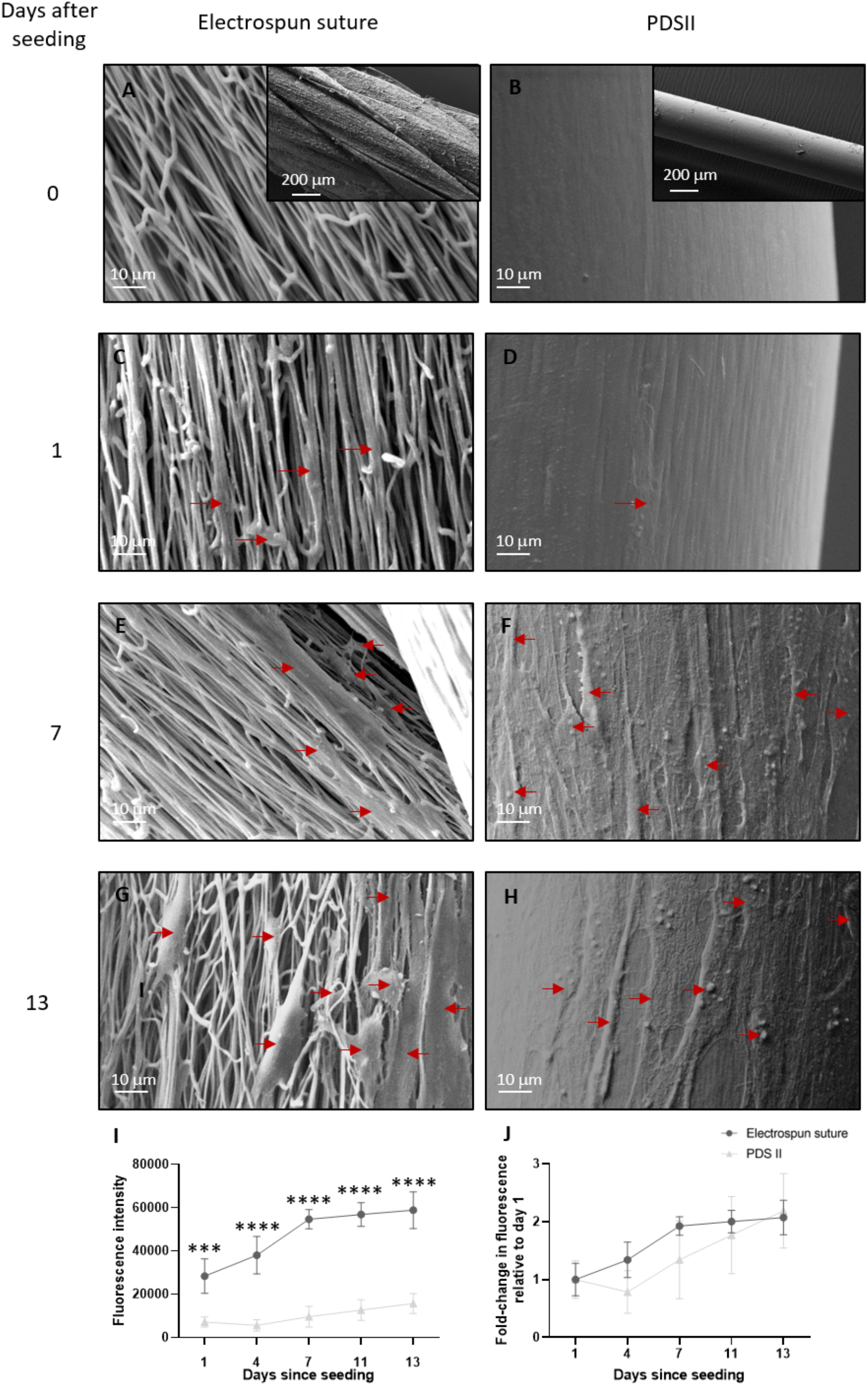
Effect of suture material on cell attachment and viability after 13 days of healthy tendon-derived stromal cell culture. (A-H) SEM images of suture materials at different points during cell culture with healthy human tendon-derived stromal cells (magnification 2000X). Red arrows point to attached cells. (A) Electrospun suture pre-seeding (x200 inset), and at days 1 (C), 7 (E), 13 (G) after seeding. (B) PDS II pre-seeding, and at days 1 (D) (x200 inset), 7 (F), 13 (H) after seeding. (I) Fluorescence intensity, indicating the number of viable cells in the sample, plotted over a period of 13 days. There was a significantly higher fluorescence intensity from the electrospun suture compared to PDS II at each time point. (J) Fluorescence was also plotted relative to day 1 to indicate proliferation independent of initial cell attachment. No statistically significant differences were seen between electrospun suture and PDS II. (I-J) represent the average results of n=4 patient samples cultured on n=3 intra-experimental replicates of each suture. Error bars indicate standard deviation.

### Electrospun suture elicits a stronger transcriptional response in tendon-derived stromal cells than PDS II

The transcriptome of tendon-derived stromal cells at baseline and cultured for 14 days on the sutures or TCP was evaluated using bulk RNA-Seq. Differentially expressed genes (Padj<0.05, LFC±1) of cells cultured for 13 days on electrospun sutures, PDS II, and TCP compared to baseline are shown using volcano plots (Figure 2). After 14 days’ culture, 1849, 667 and 61 genes were differentially expressed on electrospun sutures, PDS II and TCP, respectively, when compared to baseline (Figure 2A-C), When directly comparing the transcriptome of cells cultured on electrospun and PDS II sutures on day 13, 122 genes were differentially expressed (Figure 2D).

**Figure 2.**
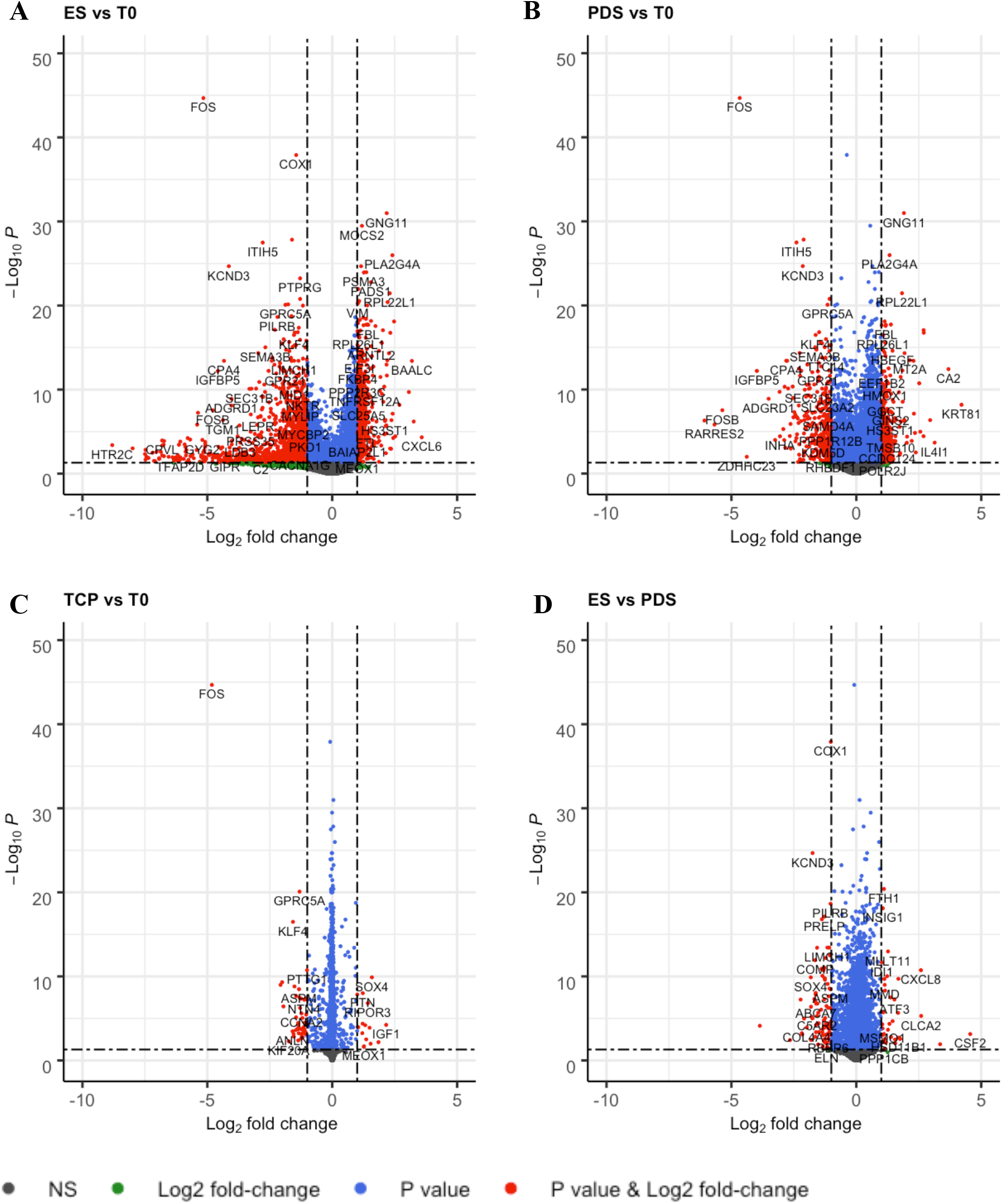
Differentially expressed genes of tendon-derived stromal cells cultured on (A) electrospun (ES) sutures, (B) PDS II sutures, or (C) tissue culture plastic control for 14 days compared to baseline control (T0). (D) Differentially expressed genes of cells cultured on either ES or PDS II for 14 days. Genes meeting the statistical significance (p<0.05) and have a log2 fold change of at least ±1 are shown in red, n=4 patients.

Hallmark gene-set enrichment analysis was used to analyse if the differentially expressed genes significantly contributed to changes in 50 well-defined biological processes (Figure 3). While both sutures upregulated expression of gene-sets associated with cell cycle progression (MYC- and E2F-Targets) and DNA repair, the electrospun suture lead to more pronounced changes than PDS II. Additional differences between the two sutures could be seen in pathways related to inflammation (IL6-JAK-STAT3 Signalling and TNFα Signalling via NF-κB) and hypoxia, which were significantly downregulated by PDS II, which is similar to TCP but not the electrospun suture. When directly comparing the two sutures, 18 gene-sets were differentially regulated (Figure 3.B) including enrichment for Mtorc 1 signalling and Myc targets, and a relative reduction in the genesets belonging to the epithelial to mesenchymal transition pathway in cells cultured on electrospun suture compared to PDS II sutures. Gene ontology gene-set enrichment analysis found that differentially regulated genes between electrospun and PDS II sutures contributed to changes in biological adhesion, cellular component morphogenesis, and (collagen containing) extracellular matrix.

**Figure 3.**
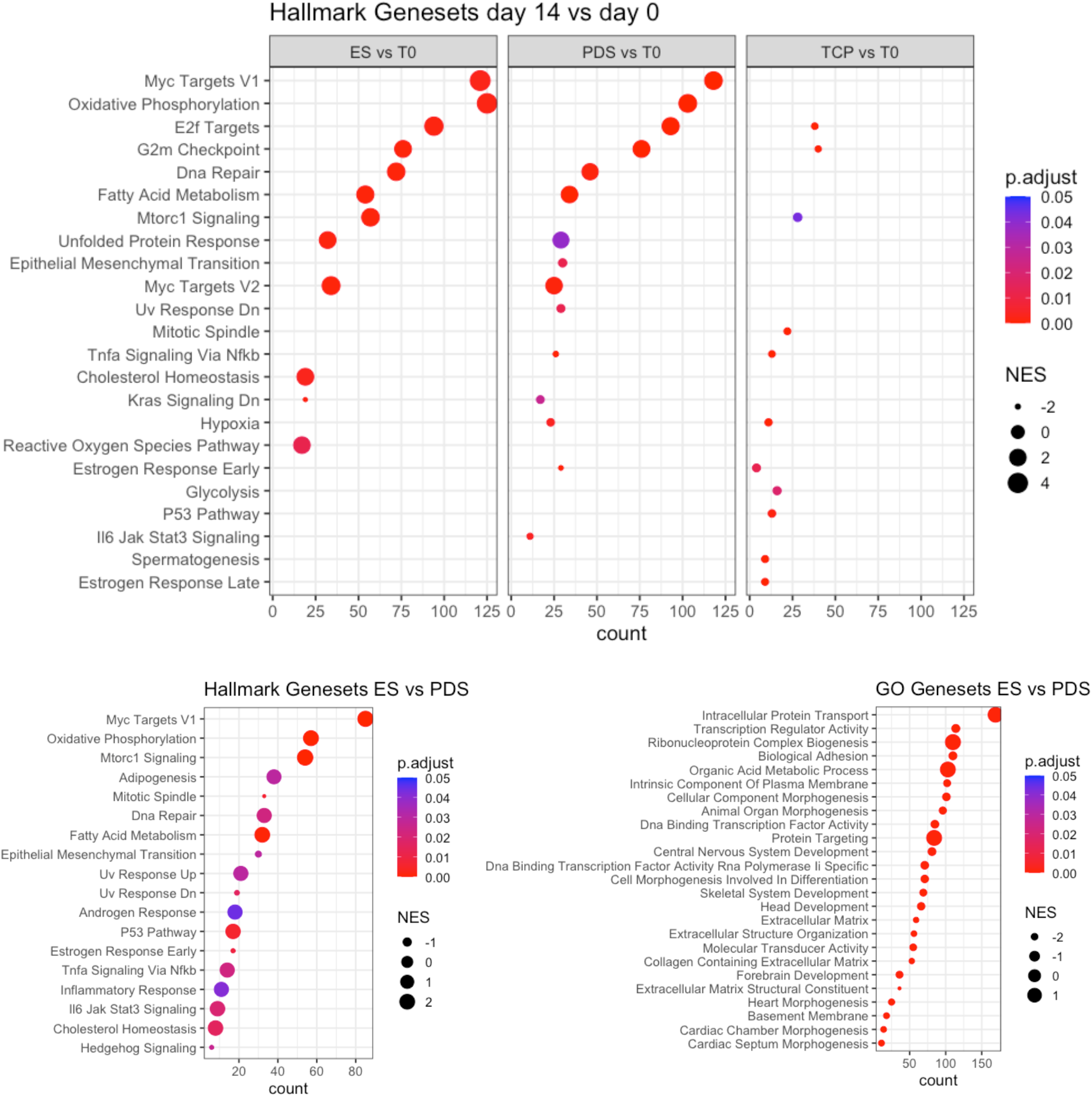
Changes in Hallmark or Gene Ontology (GO) gene sets based on differentially expressed genes. Differentially expressed genes of tendon-derived stromal cells cultured on electrospun (ES) sutures, PDS II sutures, or tissue culture plastic (TCP) control significantly contributed to changes in well-defined biological processes. Enrichment analysis of gene clusters after 14 days of cell culture on the materials was compared to baseline control (T0), and gene clusters after culture on ES and PDS II were compared directly. NES = normalised enrichment score.

### Local surface chemistry of electrospun sutures and PDS II is similar

While electrospun and PDS II sutures are both made of PDO, these results show that tendon-derived stromal cells cultured on these sutures have significant differences in attachment and transcriptional response. Differences in chemical functional groups at the suture surface may mediate altered serum protein attachment, leading to the observed differences in tendon-derived stromal cell response electrospun and PDS II sutures^24^. To establish whether differences in the structure or manufacturing processes of the sutures had resulted in differences in surface chemistry, XPS analysis was used to determine the functional groups present on the surface of the suture. Both sutures are made from PDO, containing C-C/C-H, C-O and C=O functional groups, and these groups were all present on the surface of both sutures. Although there were subtle alterations in the abundance of C-O groups on the surface of electrospun compared to PDS II sutures this did not reach statistical significance (Figure 4), suggesting that differences in structure and manufacturing processes do not strongly affect suture surface chemistry.

**Figure 4.**
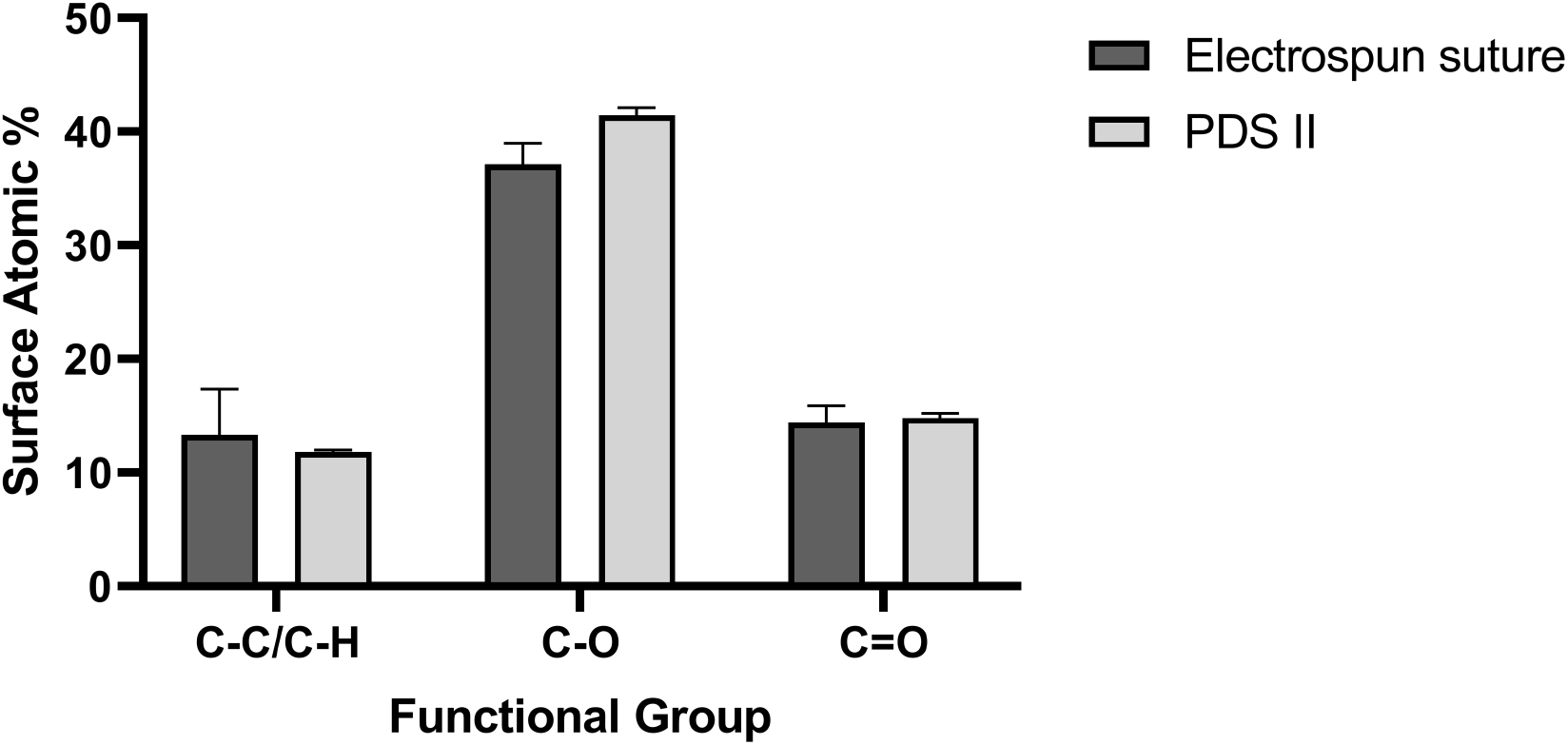
Suture surface characterisation using X-ray Photoelectron Spectroscopy. High-resolution carbon and oxygen XPS spectra, comparing carbon in various chemical states at the surface of the sutures (Surface Atomic %). No statistically significant differences in suture surface chemistry were observed. The figure represents the average results of n=3 points on a suture sample. Graphs show mean+SD.

## Discussion

Tendon disease is common and surgical repair of torn tendons is prone to failure. Electrospun materials that mimic the hierarchical structure of tendon tissues could be used to support endogenous tissue repair. This work aimed to investigate the potential of a twisted electrospun suture in the surgical repair of tendon tears. Tendon-derived stromal cells showed increased attachment to electrospun sutures. We also demonstrated that electrospun sutures induced a distinct and stronger tendon-derived stromal cell transcriptomic response when compared to PDS II sutures.

Tendon-derived stromal cells attached to and proliferated on both electrospun sutures and PDS II, but initial cell attachment to electrospun sutures was significantly higher. There were no statistically significant differences in the sutures’ surface chemistry, which would have meant similar serum protein attachment and subsequent cell attachment. However, it is likely that the greater cell attachment to electrospun suture was caused by its highly-textured surface and high surface area, compared to the smoother surface of PDS II. Indeed, electrospun sutures are composed of multiple twisted fibres with diameters that not only resemble collagen fibrils but have also been shown to promote fibroblast adhesion and infiltration^25,26^. The attachment of greater cell numbers could potentially lead to relative increases in ECM production, possibly accelerating tendon repair. Electrospun sutures have previously been shown to promote cellular infiltration *in vivo* and improved tissue integration which could reduce the rate of suture pull through^7^.

To investigate the sutures’ effects on gene expression, RNA-Seq was performed on healthy tendon-derived stromal cells after 14 days’ culture on TCP, electrospun and PDS II sutures. Ideally, sutures should stimulate a gene expression profile indicative of wound healing, upregulating pathways associated with cell proliferation and repair, without uncontrolled or sustained upregulation of fibrotic or inflammatory pathways^8 27^. Tendon-derived stromal cells cultured on electrospun suture and PDS II both upregulated gene-sets associated with the cell cycle, indicating enhanced cellular proliferation, and upregulated mTORC1 signalling, indicating upregulation of pathways relating to wound healing, protein synthesis and tendon maturation^28^. These results were more pronounced for electrospun sutures. PDS II is regarded as an immune-compatible suture, based on favourable cellular inflammatory responses in rodent models of soft tissue repair, and therefore the similarities in response to PDS II and electrospun sutures supports exploration of electrospun sutures for tendon repair^29–31^. Tendon-derived stromal cells cultured on electrospun sutures also downregulated epithelial mesenchymal transition and extracellular matrix genesets, and upregulated NF-kB genesets when compared to PDS II. This suggests they induce a wound healing response that is not strongly fibrotic, potentially minimising formation of weak scar tissue which may lead to tendon repair failure^32^. Surface chemistry, porosity and topography are all able to regulate fibroblast behaviour and may have contributed to the differences in transcriptional profile of tendon-derived stromal cells cultured on PDS II and electrospun sutures. Few genes were differentially expressed after 14 days of culture on TCP, indicating that PDS II and electrospun sutures provided cell-instructive cues and that gene expression changes on these sutures were not due to temporal changes due to prolonged culture alone. Although PDS II is considered biocompatible, it was not designed for repairing damaged tendon tissue. By allowing increased tendon-derived stromal cell attachment whilst not inducing a fibrotic transcriptional response, electrospun sutures could therefore improve the outcomes of surgical tendon repair when compared to currently used sutures.

## Perspectives

### Future directions and limitations

This work has a number of limitations. Tendon-derived stromal cells from diseased tendons would better recapitulate the response of pathological tissue to the materials. Tissue from massive rotator cuff tears (>5cm) are in greatest need of improved augmentation materials, as they have the highest rate of failure^33^. Finally, during surgical repair, the damaged tendon is rapidly infiltrated by macrophages^34^, which could alter the cellular environment that biomaterials are exposed to^35,36^. Tendon-derived stromal cells co-cultured with monocyte-derived macrophages would more accurately recapitulate the diseased tendon niche.

As Bulk-RNAseq can mask the response of cell subsets, it is also necessary to explore cell type-specific responses to various sutures. Recent reports have used single-cell RNA-Seq of healthy and diseased tendon to identify 8 or more subpopulations of tendon cells, including 5 distinct types of tendon-derived stromal cells^37,38^.

Endogenous tendon repair lasts longer than the 14 days of tissue culture conducted in this study and occurs within a loading environment^39^. However it should be noted that Rashid *et al.* examined the *in vivo* response of English Mule sheep tendon 3 months after surgical repair with a similar electrospun suture to the one described in this paper^7^. There was little inflammation and extensive cellular infiltration into the suture upon histological examination.

## Conclusions

This study compared the tendon-derived stromal cell response to clinically-used PDS II and a novel twisted PDO electrospun suture. Compared to PDS II, a currently used and safe suture, electrospun sutures demonstrated greater cell attachment and tendon-derived stromal cell transcriptomic response indicative of cell proliferation and wound healing without significant fibrosis. These results indicate that electrospun suture is a promising material that may improve the outcomes of surgical tendon repair.

## Acknowledgements

We would like to thank the NIHR Oxford Biomedical Research Centre for funding this work. We also thank the Oxford Musculoskeletal Biobank, Louise Appleton, Debra Beazley, Kim Wheway, and Bridget Watkins for collection of human hamstring tendons. We would also like to thank Hayley Morris, Joana Martins, and Antonina Lach for their guidance and assistance in manufacture of electrospun sutures. The X-ray photoelectron (XPS) data collection was performed at the EPSRC National Facility for XPS (“HarwellXPS”), operated by Cardiff University and UCL, under Contract No. PR16195. AJC receives grant funding from the NIHR, the Wellcome Trust, the MRC/UKRI and Versus Arthritis.

