## Supplementum for "*In vitro* evaluation of the response of human tendon-derived stromal cells to a novel electrospun suture for tendon repair"

S1 – Manufacture of electrospun sutures

Electrospun monofilaments of PDO were produced according to previously described methods[40] Polydioxanone (PDO, Riverpoint Medical, Portland, USA) was dissolved in 1,1,1,3,3,3-hexafluoroisopropanol (HFIP, Halocarbon Product Corporation, Atlanta, USA) at a concentration of 7% (w/v) in the presence of pyridine. Electrospinning was performed in a glove box using a single nozzle high voltage power supply system (30 kV, SL30P30/230, Spellman, West Sussex, UK) and a syringe pump (World Precision Instruments Limited, Florida, US). Filaments were produced by using a thin stainless steel wire (100 µm in diameter, Goodfellow, Huntingdon, UK) as a collector. The wire was moved underneath the nozzle at a speed of 0.5 mm/s with a bespoke winding unit. Electrospun filaments were then continuously wound up onto a motorised filament spool, separating them from the wire collector. They were then stretched manually to 3x their original length. Then, 5x50 cm stretched monofilaments were taped at one end parallel to each other, and their other end was taped to a flathead screwdriver. 300 turns/meter were performed in the S twist direction to create a ply yarn. 7 ply yarns were taped down and taped to a screwdriver as before and 150 turns/meter were performed in the Z twist direction to create electrospun sutures. The opposite twist directions make the material "twist-neutral", increasing its stability. The sutures were wound tightly around a spool, taped down and annealed for 3 hr at 65°C in a lab oven (Genlab, Widnes, UK). The samples stored in a desiccator (Bohlender, Grünsfeld, Germany) when not being handled to protect it from humidity. Electrospun sutures were cut into 2 cm pieces and melted at both ends to form suture mats that could be easily moved between wells (Fig.S1).

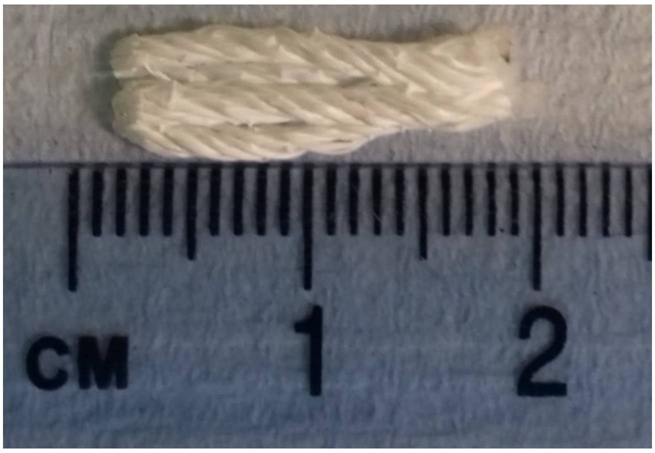


**Figure S1: Three 2 cm electrospun suture pieces were melted at both ends to form suture mats**

S2 – Tendon-derived stromal cell collection and tissue culture

Human tissue was collected with informed consent, as per the Declaration of Helsinki, with ethical approval from the Oxford Musculoskeletal Biobank (09/H0606/11 and 19/SC/0134), according to National and Institutional requirements. Biopsies of healthy waste tendon tissue from patients with Anterior Cruciate Ligament (ACL) rupture were collected during ACL reconstruction surgery, immediately transferred into sterile tubes containing D10 media and stored in a cold room when not being handled. Biopsies were prepared within 2-6 hrs by cutting into small explants under sterile conditions, using a forceps and scalpel to minimise compression of the tendon substance. 100 µL of D50 (DMEM-F12 containing 50% FCS and Penicillin-Streptomycin) culture media was added dropwise to each piece, followed by an additional 2 ml of media after 24 h. Media was changed every 3 days. After the tendon-derived stromal cells were allowed to migrate out of the explants for 7-10 days, the explants were removed. Tendon-derived stromal cells reached 80% confluence, and were then frozen in 90% FCS and 10% DMSO until required.

Tendon-derived stromal cells were thawed in warmed D10 media, centrifuged at 400 xg for 4 min and re-suspended in 20 ml media. Cells were cultured in 10 cm culture dishes (greiner bio-one, Kremsmünster, Austria) in 10 ml of D10 media. The cells were stored in an incubator (BINDER, Bohemia, NY, USA) at 37°C and 5.3% CO2. Media was changed every 3 days. Once 80% confluence was reached the cells were split. Old media was removed, and dead cells were washed away with 4 ml of D10 media. Cells were detached in 2 ml of media using a cell scraper (greiner bio-one). Scraping was performed twice, and the cells were spun down for 5 min at 0.4 rcf. The pellet was re-suspended in 1 ml of media and moved to new culture dishes. Cells were seeded at passage 3 to prevent the phenotypic drift observed at high passages[41].

### S3 – Preparing sutures for Scanning Electron Microscopy

Suture pieces were fixed for 10 min in 2.5 % glutaraldehyde (Sigma-Aldrich, St. Louis, Missouri, USA), rinsed twice in deionised water and dehydrated in a graded series of ethanol concentrations (Sigma-Aldrich) (40 %, 70 %, 90 %, 95 %, 100 %), for 2 min each. The sutures were left for 24 h in 100 µL of hexamethyldisilazane (Alfa Aesar, Haverhill, MA, USA). The sutures were gold-coated using the SC7620 Mini Sputter Coater System (Quorum Technologies, Lewes, UK).

### S4 – Cell viability assays

10% PrestoBlue was prepared by mixing 10X PrestoBlue Cell Viability Reagent (Thermo Fisher Scientific) with medium pre-warmed to 37°C. The sutures were incubated in 2mL of 10% PrestoBlue for 2h at 37°C. An empty well with 10% PrestoBlue was used to correct for background fluorescence. Due to insufficient numbers of cells, no calibration curve was generated for the PrestoBlue assay. However, previous experiments in the lab have shown that the number of tendon-derived stromal cells seeded and cultured on the sutures within 14 days is well within the limit of linearity of this assay's dose-response curve. Changes in fluorescence could therefore be used indicate changes in cell number.

100µL samples (n=3) were taken from each of the wells and pipetted into a 96-well plate (greiner bio-one). Bubbles in the well were popped using a sterile needle (BD Microlance, Franklin Lakes, NJ, USA). Fluorescence was measured using a FLUOstar Omega Microplate Reader (BMG Labtech, Aylesbury, UK) at 544 nm excitation and 590 nm emission. For each experiment, the gain was adjusted to a well with solution from cells cultured on electrospun suture, where the highest fluorescence was expected. Resazurin in the Prestoblue Cell Viability Reagent (Thermo Fisher Scientific) is reduced by living cells to the fluorescent resorufin, so the fluorescence indicated cell viability. A mean fluorescence was calculated from the samples taken from each well (n=3). The sutures were moved into fresh medium pre-warmed to 37°C after each assay.

### S5 – RNA Extraction

The sutures were incubated in 700 µL of TRIzol (Thermo Fisher Scientific) to lyse the cells. RNA was extracted using a Direct-zol RNA MicroPrep kit (Zymo Research, Irvine, CA, USA) according to the manufacturer's protocol. RNA concentration was measured using a Nanodrop (Implen, Westlake Village, CA, USA). Samples were frozen at -80°C until RNA-Seq.

### S6 – Principle Component Analysis (PCA) of tendon-derived stromal cell gene expression

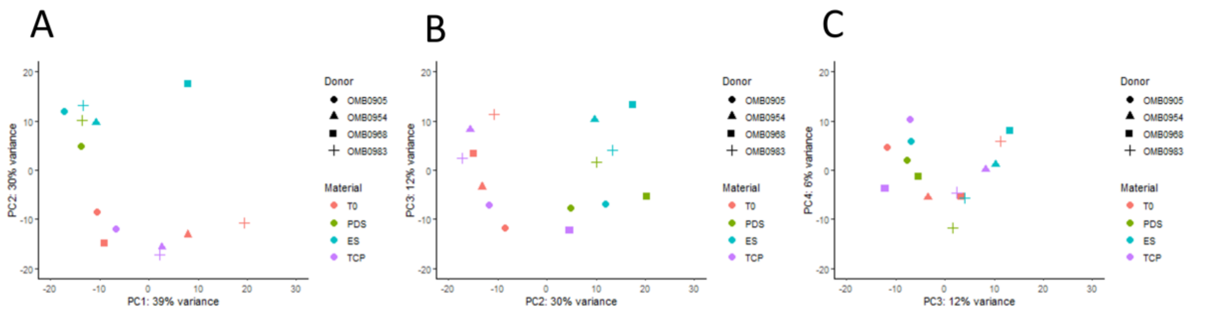

**Figure S2: Principle Component Analysis (PCA) plots of log-transformed gene expression of heathy tendon-derived stromal cells. PC2 vs PC1 (A), PC3 vs PC2 (B), and PC4 vs PC3 (C).**

Tendon-derived stromal cells were obtained from 4 donors (shown as point shapes) and gene expression was measured at time zero (T0) and after 14 days of tissue culture on 3 different materials (shown as point colours). Donor variability and material type accounted for 39% (PC1) and 30% (PC2) of variance, respectively, with TCP and T0 grouping separately to electrospun suture and PDS II. There does not appear to be differential clustering between tendon-derived stromal cells cultured on electrospun suture and PDS II up to PC4.

S7 – Analysis of count data

Log fold change shrinkage of the effect size was performed using the Apeglm package[42] to ease visualisation and ranking of genes. A Bonferroni-Hochberg Procedure was applied and an adjusted P value (Padj) was calculated to decrease the false discovery rate. Differential gene expression was visualised with volcano plots created using the EnhancedVolcano package[43]. To reveal the functional significance of the differentially-expressed genes, a Gene-Set Enrichment Analysis (GSEA) was performed genes which were differentially-expressed to a statistically significant level (Padj<0.05), using the fgsea package[44]. No fold-change cut-off was used as this is not representative of biological significance. Because the Molecular Signatures Database contains >10,000 gene sets with a great deal of redundancy, the "Hallmark Gene Set Collection"[45] was used. Normalised Enrichment Scores (NES) were calculated for these pathways. NES values describe the extent to which members of a geneset are overrepresented at the extremes of a list, which ranks these genes in order of the extent to which their expression correlates with phenotypic class distinction[46]. A Leading Edge Analysis was performed to reveal which genes are mostly responsible for the upregulation of certain biological processes.

### S8 – XPS settings

XPS data was acquired using a Kratos Axis SUPRA using monochromated Al kα (1486.69 eV) X-rays at 12 mA emission and 15 kV HT (180W) and a spot size/analysis area of 700 x 300 µm. The instrument was calibrated to gold metal Au 4f (83.95 eV) and dispersion adjusted give a BE of 932.6 eV for the Cu 2p3/2 line of metallic copper. Ag 3d5/2 line FWHM at 10 eV pass energy was 0.544 eV. Source resolution for monochromatic Al Kα X-rays is ~0.3 eV. The instrumental resolution was determined to be 0.29 eV at 10 eV pass energy using the Fermi edge of the valence band for metallic silver. Resolution with charge compensation system on <1.33 eV FWHM on PTFE. High resolution spectra were obtained using a pass energy of 20 eV, step size of 0.1 eV and sweep time of 60s, resulting in a line width of 0.696 eV for Au 4f7/2. Survey spectra were obtained using a pass energy of 160 eV. Charge neutralisation was achieved using an electron flood gun with filament current = 0.38 A, charge balance = 2 V, filament bias = 4.2 V. Successful neutralisation was adjudged by analysing the C 1s region wherein a sharp peak with no lower BE structure was obtained. Spectra have been charge corrected to the main line of the carbon 1s spectrum (adventitious carbon) set to 284.8 eV. All data was recorded at a base pressure of below 9 x 10-9 Torr and a room temperature of 294 K. Data was analysed using CasaXPS v2.3.19PR1.0. Peaks were fit with a Shirley background prior to component analysis. Quantitative data was averaged from 3 analysis spots.
